# Stick-Slip Dynamics of Migrating Cells on Viscoelastic Substrates

**DOI:** 10.1101/712059

**Authors:** Partho Sakha De, Rumi De

## Abstract

Stick-slip motion, a common phenomenon observed during crawling of cells, is found to be strongly sensitive to the substrate stiffness. Stick-slip behaviours have previously been investigated typically using purely elastic substrates. For a more realistic understanding of this phenomenon, we propose a theoretical model to study the dynamics on a viscoelastic substrate. Our model based on a reaction-diffusion framework, incorporates known important interactions such as retrograde flow of actin, myosin contractility, force dependent assembly and disassembly of focal adhesions coupled with cell-substrate interaction. We show that consideration of a viscoelastic substrate not only captures the usually observed stick-slip jumps, but also predicts the existence of an optimal substrate viscosity corresponding to maximum traction force and minimum retrograde flow which was hitherto unexplored. Moreover, our theory predicts the time evolution of individual bond force that characterizes the stick-slip patterns on soft versus stiff substrates. Our analysis also elucidates how the duration of the stick-slip cycles are affected by various cellular parameters.

## I. INTRODUCTION

Cell motility plays a key role in many important biological processes such as wound healing, morphogenesis, embryonic development, tissue regeneration to name a few [1–5]. Though motility is expressed in multiple ways, crawling happens to be the most common form of movement for eukaryotic cells. During crawling, cell forms protrusions at the leading edge which pushes the membrane forward and as a consequence the membrane exerts a backward force on the polymerising actin filaments, resulting in them ‘slipping’ rearward towards the cell center, in a process known as retrograde flow [6–8]. This process is accompanied by growth and strengthening of focal adhesions between the cell and the substrate which slow down the actin retrograde flow [9]. Thus, it allows actin polymerization to advance at the leading edge and in turn, the rate of translocation of the cell increases [1, 8, 10, 11]. The dynamic variation in retrograde flow coordinated with assembly and disassembly of focal adhesions lead to stick-slip motion.

Stick-slip is a kind of jerky motion that has been found not only in living systems but also in passive systems such as peeling of scotch tapes [12–14], earthquakes [13, 15, 16] to name a few. Stick-slip behaviour is characterised by the system spending most of it’s time in the ‘stuck’ state and comparatively a short time in the ‘slip’ state. Stick-slip dynamics has been experimentally observed on multiple occasions in crawling cells. Experiments on migrating epithelial cells showed that in the lamellipodium region the traction force decreases with increasing velocity inferring a stick-slip regime of the actin-adhesion interaction [17]. Stick slip motion has also been observed in embryonic chick forebrain neurons [18], migrating human glioma cells [19] and also in human osteosarcoma cells [20]. In case of fish keratocytes, stick-slip kind of behaviours have been found in modulation of the cell shape during crawling [21–23]. Moreover, recent experiments show that the stick-slip dynamics is strongly affected by altering the substrate stiffness. For example, a close inspection of the leading-edge motion of crawling and spreading mouse embryonic fibroblasts revealed that the periodic lamellipodial contractions are vastly substrate-dependent [24]. Further, in recent studies, cell motility was found to be maximal and actin flow rate minimal at an optimal substrate stiffness [18, 19, 25–27].

There are many theoretical studies that have contributed significantly to understand the cell migration process [28–35]. Quite a few models have also been proposed to unravel the stick-slip mechanisms. Such as, Barnhart *et. al*. developed a mechanical model to predict periodic shape change during keratocyte migration caused by alternating stick-slip motion at opposite sides of the cell trailing edge [23]. Also, leading edge dynamics, spatial distribution of actin flow, and demarkation of lamellipodium-lamellum boundary have been studied [31, 32]. Besides, simple stochastic models have provided a great deal of information on cell crawling. Stochastic bond dynamics integrated with traction stress dependent retrograde actin flow could capture the biphasic stick-slip force velocity relation [36, 37]. Another model involving stochastic linkers has also shown the presence of biphasic realtionship of traction force with substrate stiffness [38]. There are other models based on stochastic motor-clutch mechanisms which have provided many insights into substrate stiffness dependent migration process [18, 19, 27, 39, 40]. As observed in experiments, these studies reveal the existence of an optimal substrate stiffness which found to be sensitive to cell motor-clutch parameters. However, most of these studies on rigidity sensing so far are either focused on purely elastic substrate [19, 39] or on purely viscous substrate [41]; whereas, physiological extracellular matrix is viscoelastic in nature. Also, experimental studies reveal that the dynamics is greatly sensitive to the substrate viscosity [41–43]. Further, a recent study based on a modfied version of molecular clutch model and adhesion reinforcement mechanism shows that cell spreading could be affected by the substrate viscoelasticity, depending on substrate relaxation and clutch binding timescales [44].

In this paper, we present a theoretical model of the leading edge dynamics of crawling cells on a viscoelastic substrate. Our theory based on a framework of reaction-diffusion equations takes into account the retrograde flow of actin, myosin contractility, force dependent assembly and disassembly of focal adhesions integrated with cell-substrate interaction. The model predicts how these cellular components work together to give rise to the spontaneous emergence of stick-slip jumps as observed in experiments. More importantly, it elucidates the effect of variation of substrate viscoelasticity on the ‘stick-slip’ dynamics. Interestingly, it predicts, the existence of an optimal substrate viscosity corresponding to maximum traction force and minimum retrograde flow as observed in case of elastic substrate [18, 19, 39]. Our model further predicts that the cell looses its ability to differentiate between soft and stiff elastic substrate stiffness in the presence of high substrate viscosity. On the other hand, the cell can not sense the variation in substrate viscosity on a stiff substrate. Moreover, our continuum model framework captures the time evolution of individual bond force that has remained unexplored so far. These findings suggest that the nature of non-trivial force loading rate of individual bonds play a crucial role in determining the stick-slip jumps and thus explain the distinctive patterns on varying substrate compliance. Our theory also provides an analytical understanding of how the cellular parameters such as substrate stiffness, myosin activity, retrograde flow affect the duration of the stick-slip cycles.

## II. THEORETICAL MODEL

Our theory is based on the molecular clutch mechanisms proposed to describe the transmission of force from actin cytokeleton to extra cellular matrix [10, 18, 40]. The clutch module or the ‘connector’ proteins provide a dynamic link between F-actin and adhesion complexes and slow down the F-actin retrograde flow [1, 6, 8]. Our model consists of free receptors representing these ‘connector’ proteins diffusing in the actin cytoplasm. The substrate consists of a large number of adhesive ligands which can bind with these free receptors to form closed bonds as illustrated in Fig. 1. Thus, the receptors are considered to be either in free or bound states denoting open or closed ligand-receptor adhesion bonds. The ligand-receptor bonds are modelled as Hookean springs of stiffness *K*_c_. As the F-actin bundle (modelled as a rigid rod), pulled by the myosin motors [9], moves with the retrograde velocity, *v*_m_, the spring gets stretched and thus, the force on a single bond is given by multiplying the spring stiffness with the bond elongation as, *f* = *K*_c_ (*x*_b_ − *x*_sub_); where *x*_b_ is the displacement of one end of the bond attached to the actin bundle and *x*_sub_ is the displacement of the substrate (where the other end of the bond is attached). As the end of the bond attached to the actin bundle moves with the retrograde flow velocity velocity *v_m_*, *x_b_* is given by, *x_b_* = ∫ *v_m_dt*. The retrograde flow velocity of the F-actin bundle slows down with increase in the force on the closed bonds and is given by the relation, 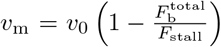 where *v*_0_ is the unloaded velocity, 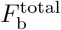 is the total traction force due to all closed bonds, and *F*_stall_ is the total force exerted by myosin motors. Here, *F*_stall_ = *n*_m_ * *F*_m_, where *n*_m_ is the number of myosin motors present and *F*_m_ is the force exerted by an individual myosin motor [9, 17, 18, 40].

**FIG. 1.**
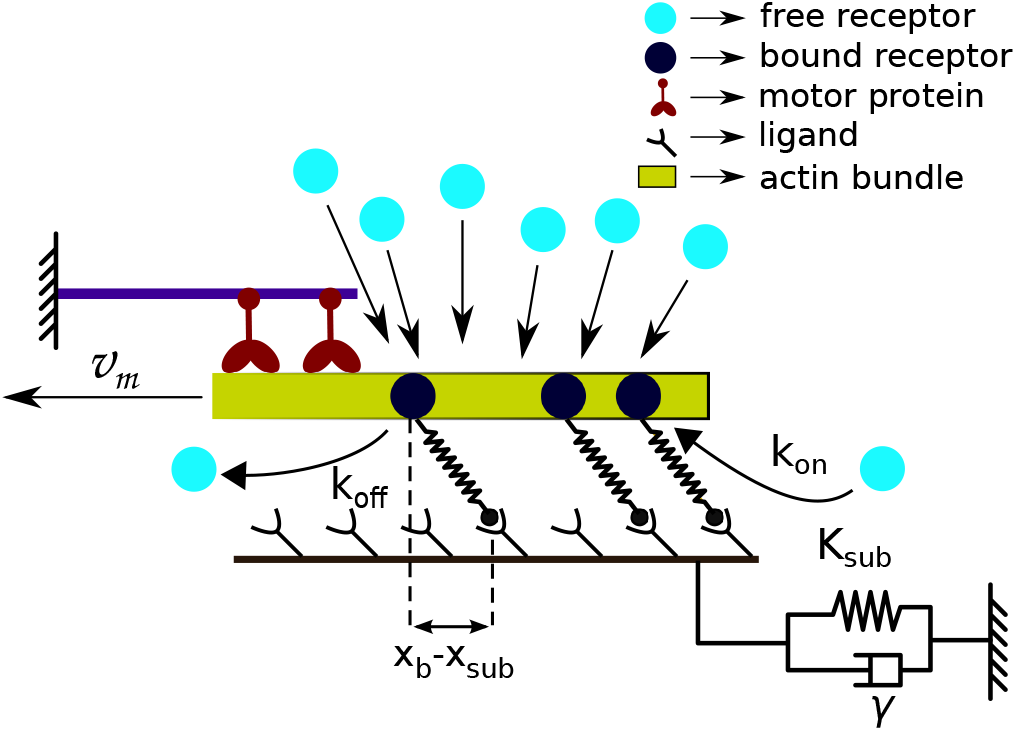
Schematic diagram of the model: free receptors (denoted by light circles) diffuse within the actin cytoplasm. The free receptors bind with ligands on the substrate to form the bound receptors (dark circles), that forms the ligand-receptor bonds. The F-actin filament is pulled by myosin motors (maroon structures) with the retrograde flow velocity, *v_m_*. Viscoelastic substrate is modelled by a spring and a dashpot. (color online)

We express the reaction between the free receptors and the ligands to form the bound receptors as

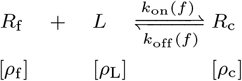

where *ρ*_f_, *ρ*_c_, and *ρ*_L_ denote the densities of free receptors, bound receptors, and ligands on the substrate respectively. The total number of free and bound receptors is taken to be conserved. The attached bound receptors do not diffuse and the influx of free receptors at the adhesion patch is assumed to be same as the outflux. Thus, 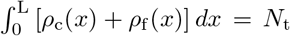 (constant). Now, the time evolution of the density of the free and the bound receptors are described by the following coupled reaction-diffusion equations,

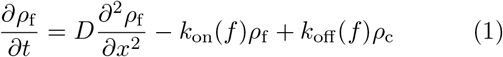

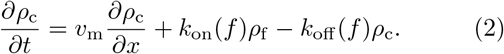

Here, the first term on the R.H.S of Eq. 1 represents the diffusion of free receptors in the actin cytoplasm. The last two terms are the reaction terms of formation of bound receptors and free receptors with respective reaction rates. For Eq. 2, the first R.H.S term denotes the drift of the bound receptors with the retrograde flow velocity, *v*_m_, as they are attached to F-actin bundle, and the other two terms are the reaction terms as in Eq. 1. In our model, motivated by the experimental findings, the association and the dissociation rates, *k*_on_(*f*) and *k*_off_(*f*), are considered to be force dependent [45–47]. We have taken, 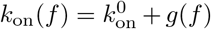; where, 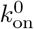 is the rate constant, and *g*(*f*) is a function of the bond force *f*; for sake of simplicity it is taken to be linear as *g*(*f*) = *ξf*. Thus, the force increases the binding rate and allows for the formation of new bonds; hence, effectively strengthening the adhesion cluster. Moreover, it has been observed that the force, upto an optimal value, helps strengthen the existing focal adhesion bonds and these force-strengthening molecular bonds are called catch bonds [48, 49]. In our model, the dissociation rate of the closed bonds is considered to demonstrate catch behavior as, *k*_off_ = *k*_s_*e*^*f*/*f*_s_^ + *k*_c_*e*^−*f*/*f*_c_^., here *k*_0_, *k*_s_, and *k*_c_ are the rate constants [50].

Moreover, in our theory, the substrate is considered to be viscoelastic in nature and has been modelled as a spring-dash pot system with spring stiffness *K*_sub_ and viscosity *γ* as shown in Fig. 1. Now, the equation of motion for the substrate is obtained by balancing the total force experienced by all the bonds with the sum of the elastic force (*K*_sub_*x*_sub_) and the viscous drag (*γẋ*_sub_) of the substrate,

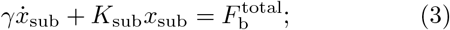

here the total traction force is given as, 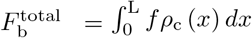.

## III. DIMENSIONLESS FORMULATION

We study the dynamics in dimensionless units. The densities have been scaled as: 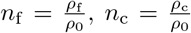, where *ρ*_0_ is the average density of the receptors and is defined as, 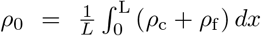. The dimensionless time is defined as *τ* = *k*_0_*t*. Thus, the dimensionless binding and unbinding rates are, 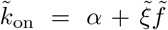 and 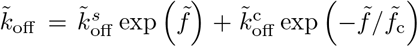, where 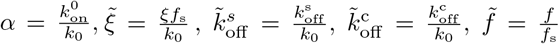 and 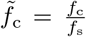. The position coordinate is scaled as 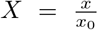, where 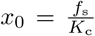. Other dimensionless variables are: 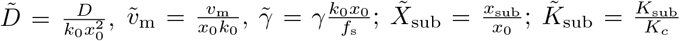; and 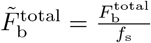.

Thus, the scaled equations of motion turn out to be,

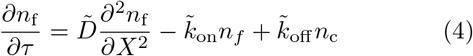

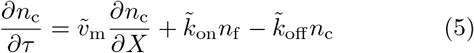

and

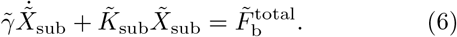

## IV. RESULTS

We have investigated the stick-slip dynamics by solving the coupled reaction-diffusion Eqs. 4-6 numerically. The equations are discretized using finite difference method on a grid of size *N* and then solved by fourth order Runge-Kutta method. The boundary conditions are taken to be such that the total number of free and bound receptors present in the system is conserved. We have studied the dynamics for a wide range of parameter values by varying system size, number of myosin motor, retrograde flow velocity, binding rates, substrate stiffness and viscosity. Here, we present the result for a representative parameter set, where values of the force dependent rate constants are kept at *α* = 2, 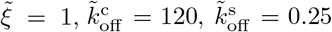 and 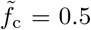; also, the unloaded velocity 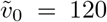, the diffusion constant 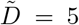, the stall force 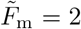, number of myosin motors *n_m_* = 100, and the system size *N* = 100. The substrate viscosity 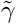 and rigidity 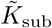 remain as variable parameters. We have also varied the diffusion constant, 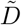; the system reaches to the steady state faster with a higher value of diffusion constant, however, the stick-slip dynamics remain qualitatively the same. Moreover, we note that even though the model parameters are scaled and dimensionless, nonetheless their values are taken from experiments, *e.g*., the dissociation rate constants are taken as 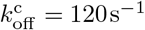 and 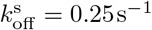 [50], whereas, unloaded myosin motor stall force is *F*_m_ = 2 pN and unloaded retrograde flow velocity is *v*_0_ = 120 nms^−1^ [18, 39]. Also, the variation of substrate elastic stiffness is considered as *K*_sub_ ~ 0.01 − 1000 pN nm^−1^ [18, 39], and the range of substrate viscosity is *γ* ~ 0.01 − 100 pN.s nm^−1^ as observed in experiments [41].

### A. Stick-slip dynamics: dependence on substrate stiffness

Figures 2 a-d show the time evolution of single bond force, total number of bonds, total traction force and corresponding retrograde flow velocity on a soft substrate. Soft substrates are very compliant and deform easily, thus, the build up of force, 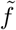, on an individual bond is also slow as shown in Fig. 2 a. Now, the increase in bond force, 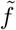, increases the binding rate, 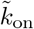, of the free receptors and at the same time, decreases the dissociation rate, 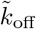, of the bound receptors upto an optimal force value due to catch bond behaviour. As a result, initially a large concentration of receptors are bound to the ligands on the substrate (shown in Fig.2 b). As the density of the bound receptors increases, they share the total traction force exerted by the substrate. Thus, the traction force slowly grows with time (Fig. 2 c) as the substrate gets deformed. The growth of traction force in turn slows down the retrograde flow of actin (Fig. 2 d). This gives rise to the ‘stuck’ state which allow actin polymerization to advance the leading edge of the cell. But as the force increases even further, the dissociation rate starts to increase. Then, the linearly growing binding rate can no longer keep up and falls below the much faster growing dissociation rate and thus, the adhesion cluster dissociates rapidly and the number of closed bonds starts decreasing. The dissociation of bonds increases the effective force on the remaining bound receptors/bonds as less number of bonds have to share the currently high value of traction force. This increases the dissociation rate even further and as a result of this feedback cycle, the bound receptors dissociate very quickly. The quick dissociation of bonds means that there is nothing to holding on or anchor to the substrate and thus, the retrograde velocity increases rapidly during what is known as the ‘slip’ state as could be seen from Fig. 2 d, and the stick-slip cycle repeats.

**FIG. 2.**
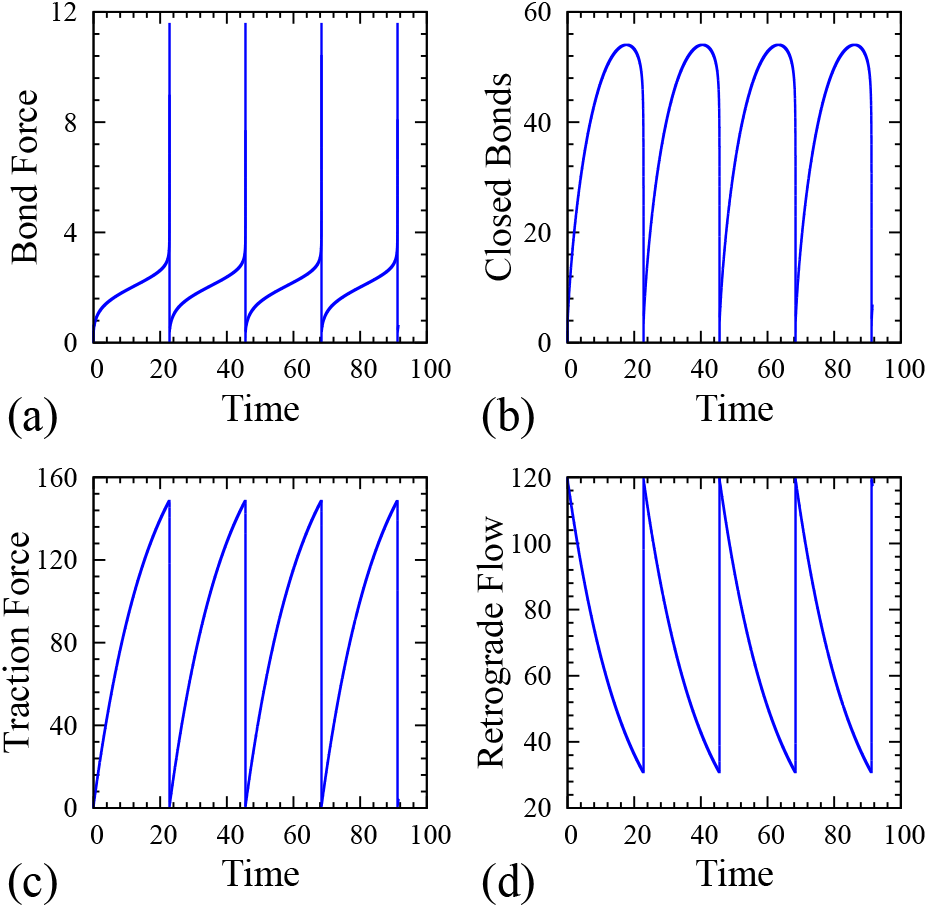
Stick-slip motion on soft substrate. (a) Time evolution of single bond force, 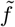. (b) time evolution of the total number of bonds, 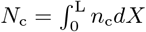. (c) corresponding evolution of the traction force, 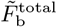, and (d) the retrograde flow velocity, 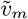, mirrors the effect of the slowing down of the actin flow with increase in the traction force. (Keeping 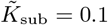 and 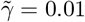, all values are dimensionless.)

On the other hand, in case of stiff substrate, as the substrate does not deform easily, force on an individual bond increases very quickly due to actin retrograde velocity as shown in Fig. 3 a. However, within this little time, due to lack of sufficient binding time, the formation of bound receptors/bonds happens to be very small as could be seen from Fig. 3 b. Now, as these small number of closed bonds are to share the total traction force exerted by the substrate, this makes the force on a single bond to increase even faster within a short time (Fig. 3 a). As a result, the exponentially varying dissociation rate dominates over linearly increasing binding rate; hence, the adhesion cluster starts dissociating even before the substrate has a substantial deformation and the traction force has attained a high value (Fig. 3 c). Lower traction force also results in a higher retrograde flow rate (shown in Fig. 3 d) as the actin filament slips backward faster and the cell thus is unable to effectively transmit forces to the stiff substrate compared to a soft substrate.

**FIG. 3.**
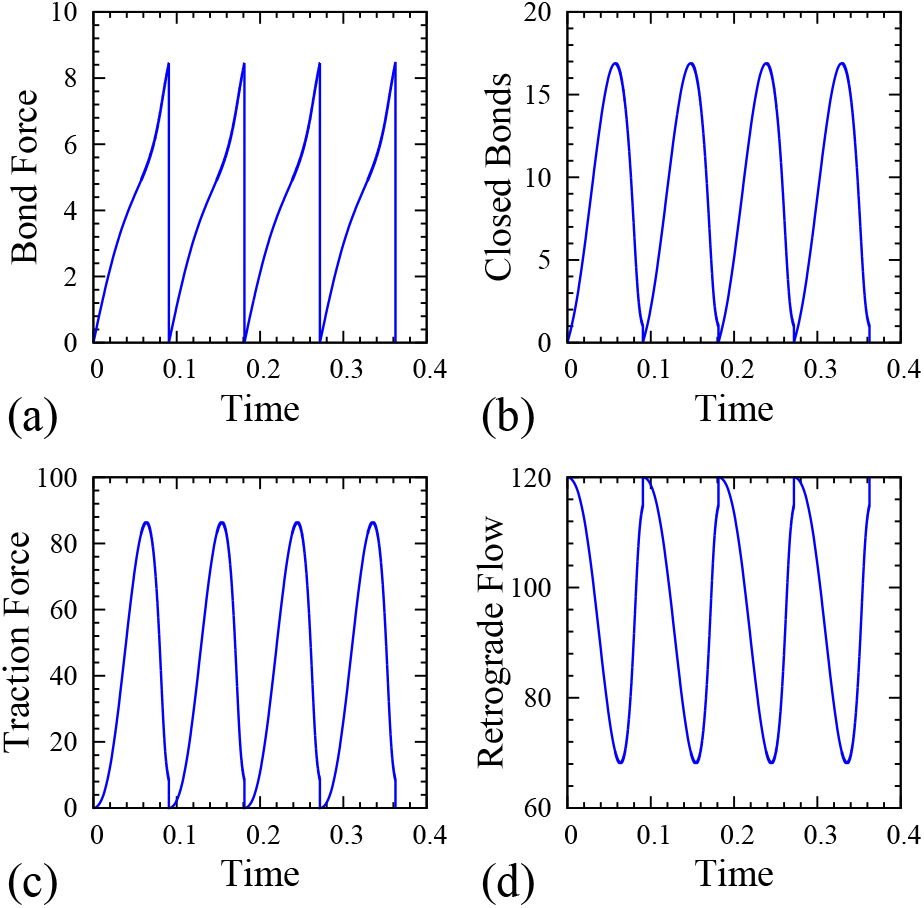
Stick-slip motion on stiff substrate. (a) Evolution of single bond force, 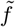, as a function of time. (b) Time evolution of the total number of bonds, 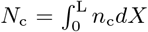, (c) corresponding evolution of the traction force, 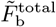, and (d) the retrograde flow velocity. (Keeping 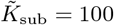 and 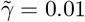)

We further investigate the stick-slip behaviours as a function of varying substrate stiffness. As observed in earlier studies [18, 19, 39], our simulations show that there exists an optimal substrate stiffness, where the mean value of the total traction force is maximum and the retrograde flow is minimum as shown in Fig. 4. This could be attributed to the difference in the nature of increase of force of an individual bond depending on the substrate stiffness as evidently seen from Fig 2 a and Fig 3 a. On a stiff substrate, since the single bond force increases rapidly, only limited number of bound receptors could form within that short time. Moreover, fast increasing force shortened the life time of the bonds; thus, resulting in lower total traction force and higher retrograde flow. However, on a softer substrate, as the substrate deforms easily, bond force increases slowly which allows for the formation of more bound receptors. Thus, the higher density of adhesion bonds results in higher value of traction force that resists the actin flow and thus, decreases retrograde velocity. As the substrate is made even more softer, the traction force and consecutively the force on individual bonds grows very slowly, due to the extreme compliance of the substrate. As a result, the system spends a large amount of time in a state of experiencing low traction force and higher retrograde velocity. This reduces the mean value of the traction force for very compliant substrates to some extent and thus increases the mean retrograde velocity.

**FIG. 4.**
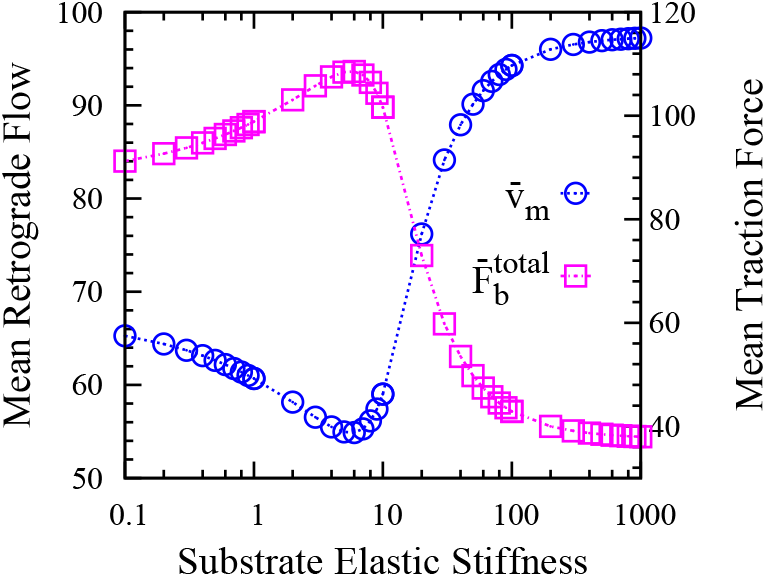
Mean value of the retrograde flow velocity 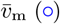 and the traction force 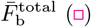 avaraged over the stick-slip cycle as a function of substrate stiffness, 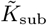 (keeping 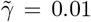 to a constant value). Average traction force is maximum and retrograde flow is minimum for an optimal value of substrate stiffness as observed in experiments [18, 19, 39].

Our theory further elucidates how the variation of substrate stiffness affects the duration of a stick-slip cycle. We obtain an analytic expression of the time evolution of the total traction force, 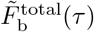 through a rudimentary calculation along with a few approximations (see supporting material). At any instant, the total traction force due to the deformation of the substrate must be balanced by summing over forces of all bound receptors/bonds. Following, the evolution of the total traction force during a stick-slip cycle (starting from a value, 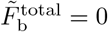 at time *τ* = 0) is given by

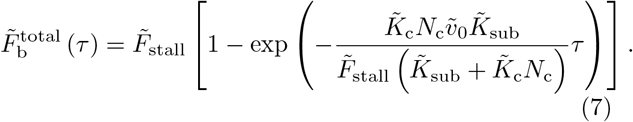

Eq. 7 can be rewritten as,

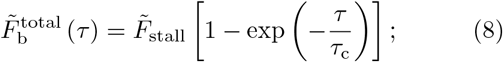

where the time constant, *τ*_c_, is of the form,

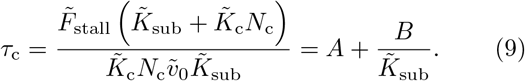

The characteristic time, *τ_c_*, is a measure of the growth rate of the total traction force and consequently, slowing down of actin retrograde flow. Therefore, it denotes the time scale corresponding to ‘stuck’ state which occupies the majority of the stick-slip cycle duration. As the slip state duration is very small compared to stuck state, the variation of *τ_c_* provides some insights into the duration of the stick-slip cycle on various system parameters, namely, substrate stiffness, bond stiffness, retrograde velocity, myosin activity, and the system size as described by Eq. 9. Moreover, we also numerically compute the duration of the stick-slip cycle as a function of substrate stiffness, 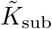, as shown in Fig. 5. Our theoretical prediction (approximated by Eq. 9) matches qualitatively well with the stick-slip cycle duration obtained from the simulation results, as seen from Fig. 5. However, a slight deviation in the stick-slip duration that is observed could be attributed to the dependence on various factors such as the reaction rates which determine the number of adhesion bonds, that in turn determines the efficiency of transmission of force from myosin motors to the substrate which have not been taken into account in our analytical prediction.

**FIG. 5.**
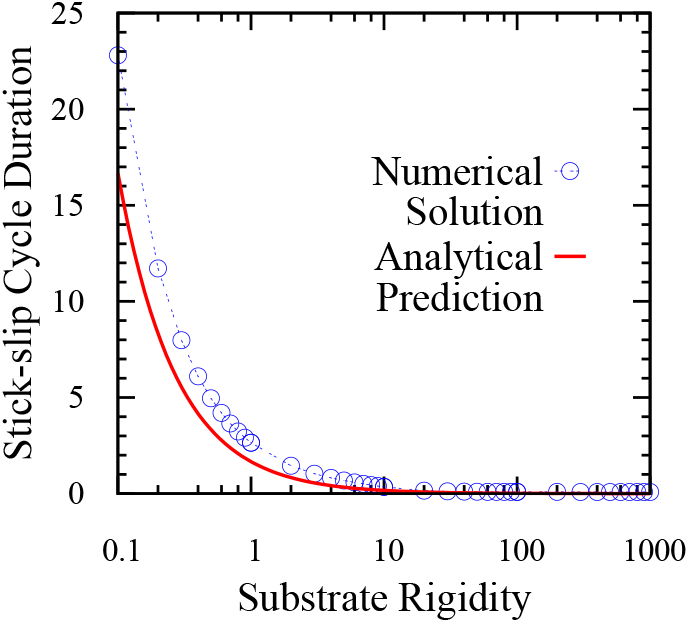
Duration of the stick-slip cycle as a function of substrate stiffness, 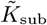. Numerical results compared with the analytical prediction of the cycle duration, given by the characteristic time, *τ*_c_, obtained from Eq. 9 (parameters are kept at same value as of numerical simulation.)

### B. Effect of the substrate viscosity on the dynamics

Recent experiments have shown that how cells spread, adhere, migrate or modulate their contractile activity vary with the extracellular matrix depending on whether it is elastic or viscoelastic in nature [41–44, 51, 52]. Moreover, the substrate stress relaxation is controlled by the viscosity; thus, it alters the cellular force transmission process and so the overall response of cells. Here, we investigate how the presence of substrate viscosity affects the stick-slip behaviour of crawling cells.

Our model predicts the presence of an optimal substrate viscosity, analogous to the previously observed optimum substrate elasticity as in the case of a purely elastic substrate, which corresponds to maximum traction force and minimum retrograde flow velocity. As seen from Fig. 6 a, keeping the substrate elastic stiffness, 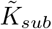 at a constant value and varying the substrate viscosity, 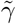, decreases the actin retrograde flow which results from increasing traction force. However, once the viscosity is increased beyond a particular optimum value, the traction force decreases and the retrograde flow increases. Viscous drag resists the deformation of the substrate, as a result of which the overall effective stiffness of the substrate increases with increasing viscosity. At a lower substrate viscosity, since the substrate relaxes faster, force on individual bonds grow slowly allowing longer interaction time with the substrate and formation of more bonds. As the adhesion cluster grows total traction force increases and hence slows down the retrograde flow. On the other hand, at higher substrate viscosity, as the substrate relaxes slowly, that causes the bond force to rise quickly without providing enough time for the formation of new bonds to share the traction force. As the bond force increases faster, it destabilizes the adhesion cluster resulting in lower traction force and higher retrograde velocity as seen from Fig. 6 a. Moreover, on variation of the substrate elasticity and observing the response of the system to change in substrate viscosity, we find that the presence of optimal viscosity and cellular sensitivity to substrate viscosity is only effective for substrates of lower elastic stiffness. Increasing substrate stiffness beyond a particular value brings about the disappearance of the optimal substrate viscosity and the cellular response becomes almost unaffected by the variation in substrate viscosity. This phenomena can be attributed to the fact that for low substrate elastic stiffness the forces on the bonds increases slowly which allows for viscosity to play it’s part in increasing the traction force and thus slowing down retrograde flow. However for a stiff substrate, the forces on the bonds are already growing fast and are soon saturated, this does not allow any time for substrate viscosity to show it’s effect; thus stripping the cell of it’s ability to differentiate variations in substrate viscosity. Moreover, it is evident from Fig. 6 b, increase in viscosity, 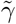, increases the effective traction force of the compliant substrate and shifts the optimal substrate stiffness for which the maximum traction occurs to a lower value 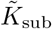. It is also interesting to note that once the substrate viscosity is increased beyond a certain threshold, the ability of the cell to differentiate between soft and rigid substrates ceases to exist. This is because at higher values of viscosity 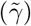, as the substrate relaxes slowly, overall stiffness effectively increases and thus, cell can not distinguish between substrates of low and high elastic stiffness.

**FIG. 6.**
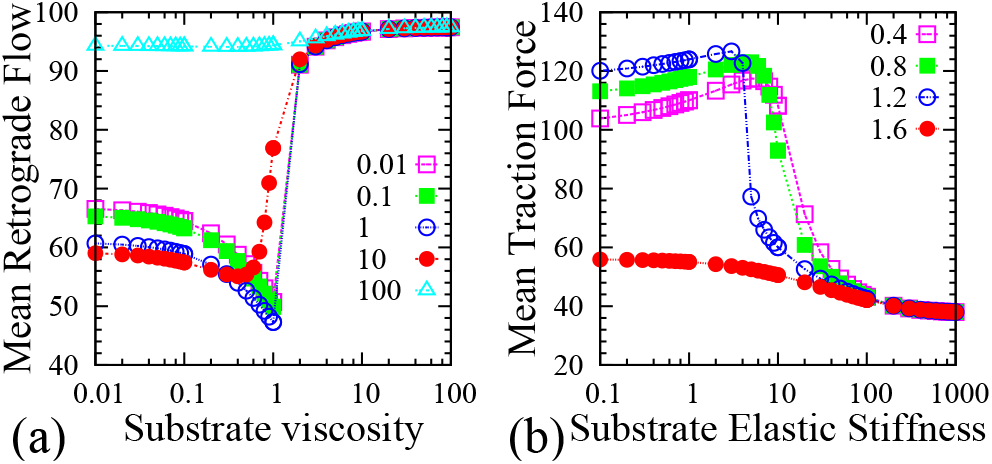
(a) Effect of mean retrograde velocity on varying substrate viscosity keeping the elastic stiffness, 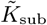, at a fixed value for different values of elastic stiffness: 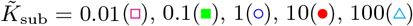. (b) Variation of mean traction force, 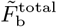, as a function of substrate stiffness, 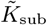, for different values of substrate viscosity: 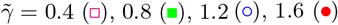

### C. Shifting of optimal stiffness

We now study how the system parameters such as binding rates, size of the adhesion patch, number of myosin motors affect the optimal substrate viscosity and also compare with that of elastic substrate. It is observed that to exhibit stick-slip behaviour, the pulling force on the F-actin bundle by the myosin motors must be balanced by the total force of the ligand-receptor bonds. However, if the total number of available receptors is too less compared to the number of myosin motors, then it results in perpetually ‘slipping’ mode, with F-actin filament moving at near it’s unloaded velocity, *v*_0_. The reverse scenario can also take place where the number of bonds is much higher compared to the number of myosin motors, so that it slows down the retrograde velocity to zero, thus resulting in a permanently ‘stuck’ state. However, if the number of myosin motors and the number of bonds, *i.e*., the size of the ligand-receptor adhesive patch are varied appropriately, the stick-slip dynamics is restored.

In our simulation, the simultaneous change in number of motors and the size of the adhesive patch is taken into consideration by changing the grid size *N* and also modifying *F*_stall_ = *n*_m_ * *F*_m_, where *n*_m_ denotes the number of myosin motors pulling the F-actin filament. Fig 7 a and Fig 7 b show the mean retrograde flow for varying system size *N* and *n*_m_; where in (a), it is plotted as a function of substrate elastic stiffness, 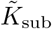, keeping the viscosity, 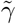, at a constant value and in (b), as a function of substrate viscosity, 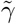, keeping 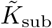 constant. As seen from Fig. 7 a, with increase in the system size, as more number of receptors/bonds can bind with the ligand on the substrate, the total bond force increases which balances the traction force by the substrate; thus, the optimal elastic stiffness for minimum retrograde flow shifts to a higher value. Fig. 7 b shows that increase in the system size also brings similar shift in the optimal substrate viscosity towards a higher value. This is because, higher viscosity results in slow stress relaxation, thus, increases the effective combined stiffness of the substrate and allows transmission of larger traction force as observed in case of pure elastic stiffness.

**FIG. 7.**
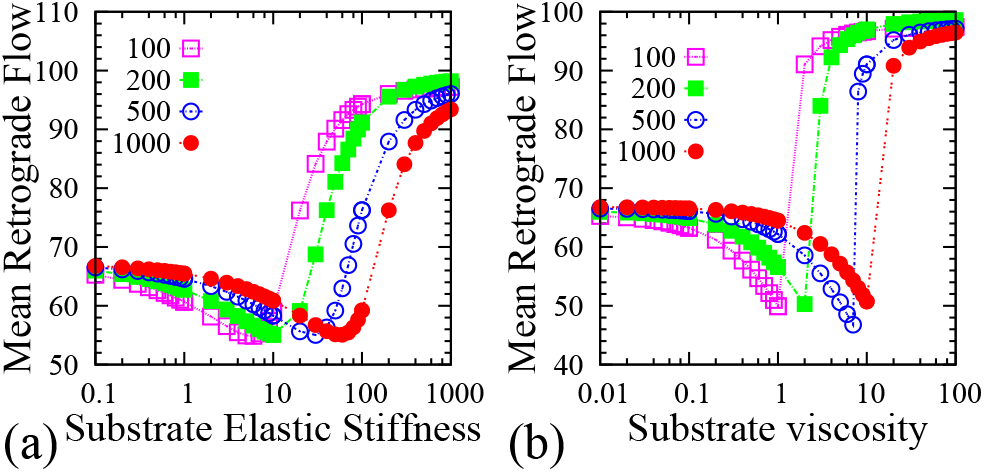
Variation in the total number of myosin motors and the size of the adhesion cluster causes shift in (a) the optimal substrate stiffness and also in (b) the optimal viscosity. (Number of motors, *n*_m_ and total number of receptors, *N*_t_ have been varied equally.) 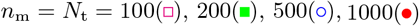

We have, further, compared the results for varying binding rates and also studied without the force induced adhesion reinforcement. It is found that the force dependent rate provides an added flexibility for changing the optimal substrate stiffness or the optimal viscosity to match the experimentally observed values for different cell types. This is achieved because changing the force induced rate results in variation of the total number of closed bonds, thus, changes the total traction force and the retrograde flow velocity and consequently shifts the optimal stiffness.

## V. DISCUSSIONS

We have developed a theoretical model to study the ‘stick-slip’ motion at the leading edge of a crawling cell. The extracellular matrix has been modelled as a viscoelastic system to better mimic the biological substrates, as opposed to the generally modelled pure elastic substrates. Our continuum model framework comprising of coupled reaction-diffusion equations predicts the time evolution of force on an individual bond during a stick-slip cycle, that could not be captured in existing stochastic model frameworks. Our study reveals that the loading rate of single bond force is distinctively different on soft substrate compared to stiff substrate. It plays a crucial role in determining the pattern of the stick-slip jumps on varying substrate rigidity. It is also worth noting that our continuum model description reduces the computational time required for averaging of the dynamical quantities as compared to stochastic models where the averaging needs to be done over a large number of trajectories to extract useful statistical information. It may also be mentioned that in our mean field model, for simplicity, we have considered that all bonds experience a homogeneous force. To check the validity of this assumption, we have also simulated a stochastic analogue to our model by taking into account the inhomogeneity of bond forces. We have observed that there is no significant difference in the overall dynamics when averaged over a large statistics it remains qualitatively the same. Further, in our model, motivated by the experimental findings, the bond association and dissociation rates are considered to be force dependent. Experimentally observed force induced reinforcement of adhesion complexes has been incorporated in the binding rate as well as through the ‘catch bond’ behaviour of adhesion complexes [48–50]. Our analysis also elucidates the dependence of the duration of the stick-slip cycle on various cellular parameters, for example, how it is affected by myosin activity, retrograde flow, or substrate stiffness. Our theory further suggests that the viscoelasticity of the substrate plays a central role in driving the cell migration process. In our model, viscoelasticity of the substrate has been incorporated by a standard Kelvin-Voigt model; however, one could also model it by a standard linear viscoelastic material as considered by Gong et al [44]. Importantly, our model reveals the existence of an ‘optimal’ substrate viscosity where the traction force is maximum and the retrograde flow is minimum similar to the variation of elastic substrate stiffness. It also predicts that the cell can sense the effect of substrate viscosity only when the substrate stiffness is low. The optimum in substrate viscosity disappears when the substrate stiffness is increased. This is because on stiffer substrate the individual bonds already experience a rapid building of tension due to high stiffness giving rise to high retrograde flow velocity and thus, substrate viscosity does not have any significant effect. Interestingly, our theory shows that the cell also loses the ability to sense variations in elastic substrate stiffness in the presence of a high substrate viscosity. This is because at higher viscosity, substrate relaxes slowly and gets stiffened up, effectively similar to stiff elastic substrate that causes the bond force to build up rapidly leading to fast disintegration of adhesion bonds and thus does not allow the cell to distinguish the variation in soft versus stiff substrate elasticity. This indicates the importance of substrate stress relaxation process in cell motility. As in experiments, cell crawling has been found to be most efficient on an optimal substrate stiffness, it could further be tested by altering the viscosity or a combination of both viscosity and elasticity to see how cells respond to viscoelastic tissues and interpreting these responses can further be useful to modulate the cellular behaviour in order to fine-tune many biophysical applications such as tissue engineering, cancer research [40] and regenerative medicine etcetera.

## ACKNOWLEDGMENTS

The authors acknowledge the financial support from Science and Engineering Research Board (SERB), Grant No. SR/FTP/PS-105/2013, Department of Science and Technology (DST), India.

## Appendix A: Analytical estimation of the duration of a stick-slip cycle

Considering an elastic substrate, the traction force due to the deformation of the substrate will be given by 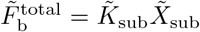, where 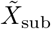 denotes the displacement of the substrate and 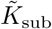 is the substrate stiffness. At any instant, the traction force must be balanced by summing over forces of all ligand-receptor bonds. Now, the total bond force could be calculated as, 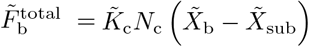, where *N*_c_ is the total number of closed bonds at any instant, 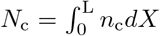, the elongation of the bond is given by 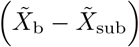. Here 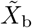 is the displacement of one end of the bond attached to the actin filament, thus, 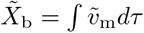. Here, 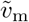 is the dimensionless retrograde flow given by 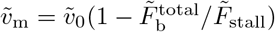.

Now, the expression for the traction force at any time *τ* can be rewritten as,

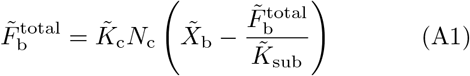

Differentiating Eq. A1 w.r.t. the dimensionless time *τ* and using the relation, 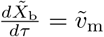, we obtain,

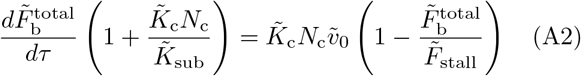

Thus, the evolution of the total traction force during a stick-slip cycle starting from a value 0 at time *τ* = 0 can be given by the solution of Eq. A2,

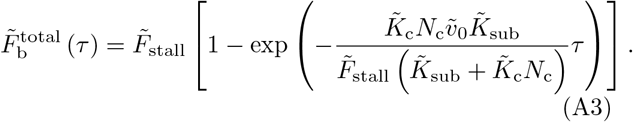

## References

[1] Gardel, M. L., Schneider, I. C., Aratyn-Schaus, Y., and C. M. Waterman. 2010. Mechanical integration of actin and adhesion dynamics in cell migration. Annu. Rev. Cell Dev. Biol. 26:315–333.

[2] Ridley, A. J., Schwartz, M. A., Burridge, K., Firtel, R. A., Ginsberg, M. H., Borisy, G., Parsons, J. T., and A. R. Horwitz. 2003. Cell migration: integrating signals from front to back. Science. 302(5651):1704–1709.

[3] Friedl, P., and D. Gilmour. 2009. Collective cell migration in morphogenesis, regeneration and cancer. Nat. Rev. Mol. Cell Biol. 10:445–457

[4] Dahlin, C., Linde, A., Gottlow, J., and S. Nyman. 1988. Healing of Bone Defects by Guided Tissue Regeneration. Plastic and Reconstructive Surgery. 81(5):672–676

[5] Krawczyk, W. S. 1971. A pattern of epidermal cell migration during wound healing. J Cell Biol. 49:247–263

[6] Ananthakrishnan, R. and A. Ehrlicher. 2007. The Forces Behind Cell Movement. Int. J. Biol. Sci. 3(5):303–317.

[7] Lin, C. H., and P. Forscher. 1995. Growth cone advance is inversely proportional to retrograde F-actin flow. Neuron. 14(4):763–771.

[8] Jurado, C., Haserick, J. R., and J. Lee. 2005. Slipping or gripping? Fluorescent speckle microscopy in fish keratocytes reveals two different mechanisms for generating a retrograde flow of actin. Mol Biol Cell. 16(2):507–518.

[9] Greenberg M.J., Arpag G., Tzel E., and E. M. Ostap. 2016. A perspective on the role of myosins as mechanosensors. Biophys. J. 110:2568–2576.

[10] Mitchison, T.J., and M. Kirschner. 1988. Cytoskeletal dynamics and nerve growth. Neuron. 1:761–772.

[11] Hu, K., Ji, L., Applegate, K. T., Danuser, G. and C. M. Waterman-Storer. 2007. Differential transmission of actin motion within focal adhesions. Science. 315:111–115

[12] De, R., Maybhate, A., and G. Ananthakrishna. 2004. Dynamics of stick-slip in peeling of an adhesive tape Phys. Rev. E. 70:046223

[13] G. Ananthakrishna and R. De. 2007. Dynamics of Stick-Slip: Some Universal and Not So Universal Features. Lecture Notes in Physics, Springer, Berlin. vol. 705:423–457

[14] De, R. and G. Ananthakrishna. 2005. Missing physics in stick-slip dynamics of a model for peeling of an adhesive tape. Phys. Rev. E vol 71:055201(R)–055204(R)

[15] De. R., Maybhate, A., and G. Ananthakrishna, 2004. Power laws, precursors and predictability during failure *Europhys*. Lett. 66(5):715–721

[16] Burridge, R. and L. Knopoff. 1967. Model and Theoretical Seismicity Bulletin of the Seismological Society of America. Vol. 57, 3:341–371

[17] Gardel, M. L., Sabass, B., Ji, L., Danuser, G., Schwarz, U. S. and C. M. Waterman. 2008. Traction stress in focal adhesions correlates biphasically with actin retrograde flow speed. J. Cell Biol. 183:999–1005

[18] Chan, C. E., and D. J. Odde. 2008. Traction dynamics of filopodia on compliant substrates. Science. 322:1687–1691.

[19] Bangasser, B.L., Shamsan, G. A., Chan, C. E., Opoku, K. N., Tuzel, E., Schlichtmann, B. W., Kasim, J. A., Fuller, B. J., McCullough, B. R., Rosenfeld, S. S. and D. J. Odde. Shifting the optimal stiffness for cell migration. Nat. Commun. 2017 8:15313

[20] Aratyn-Schaus Y and Gardel M L. 2010. Transient frictional slip between integrin and the ECM in focal adhesions under myosin II tension Curr. Biol. 20:1145–1153.

[21] Lacayo, C. I., Pincus, Z., VanDuijn, M. M., Wilson, C. A., Fletcher, D. A., Gertler, F. B., Mogilner, A. and J. A. Theriot. 2007. Emergence of large-scale cell morphology and movement from local actin filament growth dynamics. PLoS Biol. 5:e233.

[22] Lee, J., Ishihara, A., Theriot, J. and K. Jacobson. 1993. Principles of locomotion for simple-shaped cells. Nature.; 362, 167–171.

[23] Barnhart,E. L., Allen, G. M., Julicher, F. and J. A. Theriot. 2010. Bipedal Locomotion in Crawling Cells. Biophys. J. 98:933–942.

[24] Giannone, G., Dubin-Thaler, B. J., Doebereiner, H. G., Kieffer, N., Bresnick, A. R. and M. P. Sheetz. 2004. Periodic lamellipodial contractions correlate with rearward actin waves. Cell. 116:431–443.

[25] Stroka, K. M., and H. Aranda-Espinoza. 2009. Neutrophils display biphasic relationship between migration and substrate stiffness. Cell Motil. Cytoskeleton. 66:328–341.

[26] Peyton, S. R., and A. J. Putnam. 2005. Extracellular matrix rigidity governs smooth muscle cell motility in a biphasic fashion. J. Cell. Physiol. 204:198–209.

[27] Elosegui-Artola, A., Trepat, X. and P. Roca-Cusachs. 2018. Control of Mechanotransduction by Molecular Clutch Dynamics. Trends in Cell Biology. 28:356–367

[28] Rangarajan, R., and M. H. Zaman. 2008. Modeling cell migration in 3D: status and challenges. Cell Adhes. Migr. 2:106–109.

[29] DiMilla, P. A., Barbee, K. and D. A. Lauffenburger. 1991. Mathematical model for the effects of adhesion and mechanics on cell migration speed. Biophys. J. 60:15–37.

[30] Danuser, G., Allard, J., and A. Mogilner 2013. Mathematical modeling of eukaryotic cell migration: insights beyond experiments. Annual Review of Cell and Developmental Biology. 29:501–528.

[31] Shemesh, T., Bershadsky A. D., and M. M. Kozlov. 2012. Physical model for self-organization of actin cytoskeleton and adhesion complexes at the cell front. Biophys. J. 102:1746–1756.

[32] Craig, E. M., Stricker, J., Gardel, M. and A. Mogilner. 2015. Model for adhesion clutch explains biphasic relationship between actin flow and traction at the cell leading edge. Phys. Biol. 12:035002

[33] Kabaso, D., Shlomovitz, R., Schloen, K., Stradal, T. and N. S. Gov, 2011. Theoretical model for cellular shapes driven by protrusive and adhesive forces. PLoS Comput. Biol. 7:e1001127.

[34] Rubinstein, B., Fournier, M. F., Jacobson, K., Verkhovsky, A. B. and A. Mogilner. 2009. Actin-myosin viscoelastic flow in the keratocyte lamellipod. Biophys. J. 97:1853–1863.

[35] Kruse, K., Joanny, J. F., Julicher, F. and J Prost. 2006. Contractility and retrograde flow in lamellipodium motion. Phys Biol. 3(2):130–137.

[36] Sabass, B. and U. S. Schwarz, 2010. Modeling cytoskeletal flow over adhesion sites: competition between stochastic bond dynamics and intracellular relaxation. J. Phys. Condens. Matter. 22:194112.

[37] Li, Y., Bhimalapuram, P. and A. R. Dinner. 2010. Model for how retrograde actin flow regulates adhesion traction stresses. J. Phys. Condens. Matter. 22:194113.

[38] P Sens. 2013. Rigidity sensing by stochastic sliding friction. Europhys Lett, 104(3):38003.

[39] Bangasser, B. L., Rosenfeld, S. S. and D. J. Odde. 2013. Determinants of maximal force transmission in a motor-clutch model of cell traction in a compliant microenvironment. Biophys. J. 105:581–592.

[40] Elosegui-Artola, A., Bazellires, E., Allen, M. D., Andreu, I., Oria, R., Sunyer, R., Gomm, J. J., Marshall, J. F., Louise Jones, J., Trepat. X and P. Roca-Cusachs. 2014. Rigidity sensing and adaptation through regulation of integrin types. Nature Mater. 13:631–637

[41] Bennett, M., Cantini, M., Reboud, J., Cooper, J. M., Roca-Cusachs, P. and M. Salmeron Sanchez. 2018. Molecular clutch drives cell response to surface viscosity, Proc. Natl. Acad. Sci. U.S.A. 115(6):1192–1197.

[42] Chaudhuri, O., Gu, L., Darnell, M., Klumpers, D., Bencherif, S. A., Weaver, J. C., Huebsch, N. and D. J. Mooney. 2015. Substrate stress relaxation regulates cell spreading. Nature Commun. 6:6364.

[43] Lautscham, L. A., Lin, C. Y., Auernheimer, V., Naumann, C.A., Goldmann, W. H. and B. Fabry, 2014. Biomembrane-mimicking lipid bilayer system as a mechanically tunable cell substrate, Biomaterials. 35:3198–3207

[44] Gong, Z., Szczesny, S. E., Caliari, S. R., Charrier, E. E., Chaudhuri, O., Cao, X., Lin, Y., Mauck, R. L., Janmey, P. A., Burdick, J. A., and Shenoy, V. B. 2018. Matching material and cellular timescales maximizes cell spreading on viscoelastic substrates. Proc. Natl Acad. Sci. U.S.A. 115, E2686–e2695 (2018).

[45] Balaban, N.Q., Schwartz, U.S., Riveline, D., Goichberg, P., Tzur, G Sabanay, I., Mahalu, D., Safran, S., Bershadsky, A., Addadi, L., and B. Geiger, 2001. Force and focal adhesion assembly: a close relationship studied using elastic micropatterned substrates. Nat. Cell Biol. 3:466–472.

[46] Tan, J. L., Tien, J., Pirone, D. M., Gray, D. S., Bhadriraju, K. and C. S. Chen. 2003. Cells lying on a bed of microneedles: an approach to isolate mechanical force. Proc. Natl Acad. Sci. U.S.A. 100:1484–1489.

[47] De, R. 2018. A general model of focal adhesion orientation dynamics in response to static and cyclic stretch, Communications Biology. 1: 81

[48] Thomas, W. E., Vogel, V. and E. Sokurenko, 2008. Biophysics of catch bonds. Annu. Rev. Biophys. 37:399–416

[49] Marshall, B. T., Long, M., Piper, J. W., Yago, T., McEver, R. P. and C. Zhu. 2003. Direct observation of catch bonds involving cell-adhesion molecules. Nature. 423:190–193.

[50] Pereverzev, Y., Prezhdo, O. V., Forero, M., Sokurenko, E. and W. Thomas. 2005. The two-pathway model for the catch-slip transition in biological adhesion. Biophys. J. 89:1446–1454

[51] De, R., Zemel, A. and S. A. Safran. 2010. Theoretical Concepts and Models of Cellular Mechanosensing. Methods Cell Biol. 98:143–175.

[52] De, R., Zemel, A. and S. A. Safran. Dynamics of cell orientation. 2007. Nature Physics. 3:655–659.

